# Meta-analysis on the effects of chemical stressors on freshwater ecosystem functions

**DOI:** 10.1101/2023.10.25.563649

**Authors:** Alexander Feckler, Ralf Schulz, Ralf B. Schäfer

## Abstract

Despite considerable progress in our predictive capacity for the response of different trophic levels to chemical stressors, generalizable relations between chemical stressors and ecosystem functions are lacking. We addressed this knowledge gap by conducting a meta-analysis (153 studies; 350 observations) on the responses of freshwater ecosystem functions (community respiration, organic matter decomposition, nutrient cycling, photosynthesis, and primary production) under controlled conditions (laboratory or outdoor mesocosms) to pesticides, pharmaceuticals, and metals. We identified monotonic dose-response relationships between standardized chemical concentrations, in terms of toxic units, for selected chemical use groups and organic matter decomposition by decomposer-detritivore-systems as well as photosynthesis by algae and macrophytes. By contrast, consistent relationships were not found for other ecosystem functions, such as organic matter decomposition by microbial decomposers alone and primary production under the conditions studied. Importantly, the shape and direction of the relationships identified here match those reported in field-based studies, indicating a decrease in functioning as chemical stress increases, strengthening the ecological relevance of our findings. Finally, we found a disconnect between regulatory ecological quality targets and ecological outcomes, highlighting a need to re-evaluate risk assessment approaches if they are supposed to be ecologically meaningful and protective of ecosystem functions.

## 1. Introduction

Freshwater habitats (streams, rivers, lakes, and ponds) host at least 9.5% of the currently described animal species biodiversity despite comprising just around 0.01% of the global water resources and covering 2.3% of the global surface area (Shiklomanov, 1993; Strayer and Dudgeon, 2010; Reid et al., 2019). Furthermore, they play a key role with respect to ecosystem functions. Approximately 20% of all land-based nitrogen sources are removed by denitrification in freshwaters (Seitzinger et al. 2006) and the transport and partial processing of carbon in streams and rivers is equivalent to about 25% of the estimated terrestrial net ecosystem production globally (Battin et al., 2023). Despite their paramount importance, freshwater habitats experience more pronounced declines in biodiversity compared to terrestrial ecosystems, leading to what has been labelled as the “freshwater crisis” (Harrison et al., 2018). The freshwater crisis is the result of an interplay of anthropogenic disturbances: over-exploitation, chemical pollution, alterations in water flow patterns, habitat destruction or degradation, and the introduction of non-native species (Reid et al., 2019). However, chemical pollution, except for nutrients, has often been marginalized in freshwater ecosystem studies, even though it can play a substantial role in deteriorating these ecosystems (Bernhardt et al., 2017). Extensive meta-analyses on continental and global scales consistently reveal that a substantial proportion of water bodies are affected by chemicals. In Europe, nearly half of the evaluated water bodies were identified as being at risk due to chronic chemical effects (Malaj et al., 2014). Similarly, on a global scale, insecticide contamination was observed to surpass thresholds indicating ecological impacts in nearly 70% of stream sites within agricultural landscapes (Stehle and Schulz, 2015). Recent investigations underscore the substantial level of chemical risk faced by freshwater organisms, both in Europe and the United States (Schulz et al., 2021; Wolfram et al., 2021).

Numerous studies showed that toxic chemicals can influence the organisms providing pivotal ecosystem functions or the functions themselves, but their results remain scattered and generalizations on relationships remain open. Meta-analyses and reviews (e.g., Crane et al., 2006; Kümmerer, 2009; Schäfer et al., 2012b) showed that both microorganisms and invertebrates, which are crucial for ecosystem functions like nutrient cycling, organic matter decomposition, and primary production, are impacted by pharmaceuticals and pesticides at environmentally relevant concentrations. Additionally, several meta-analyses and reviews have drawn connections between effects of chemical stressors on specific groups of organisms and changes in aquatic ecosystem functions these organism groups contribute to, such as organic matter decomposition (Beaumelle et al., 2020; Feckler and Bundschuh, 2020; Ferreira et al., 2016; Peters et al., 2013; Schäfer et al., 2012b), photosynthesis (Feckler et al., 2015; Kovalakova et al., 2020), primary production (Feckler et al., 2015; Guo et al., 2015; Kovalakova et al., 2020; Peters et al., 2013; Schäfer et al. 2012a), nutrient cycling (Peters et al., 2013), and ecosystem respiration (Schäfer et al. 2012a). Recent studies also demonstrated that chemicals from the same chemical type and with the same mode of toxic action (MoA) exhibit qualitatively similar effects in communities (Halstead et al., 2014; Rumschlag et al., 2019; Rumschlag et al., 2020). Thus, it can be hypothesized that the response patterns of ecosystem functions towards chemicals from the same chemical type and MoA display a generalizable relationship between the intensity of chemical stress and the rate of ecosystem function. An earlier review by Peters et al. (2013) failed to establish consistent dose-response relationships between various groups of toxic chemicals and several ecosystem functions, such as nutrient cycling, organic matter decomposition and primary production. However, these authors rather aimed at identifying effect thresholds and only considered studies that exceeded a defined negative effect size instead of the full spectrum of responses of ecosystem functions. Indeed, the responses of ecosystem functions to toxic chemicals can be negative (e.g., Beaumelle et al., 2020; Robson et al., 2020; Rosi-Marshall and Royer, 2012), positive (e.g., Corcoll et al., 2015; Richmond et al., 2019) or non-distinguishable from a control treatment or a reference (e.g., Jepsen et al., 2019; Schäfer et al., 2012a). Thus, it remains open whether an analysis of a broader number of studies, including novel studies that have been conducted in the last decade, enables the derivation of generalizable relationships between chemical stressors and ecosystem functions.

In the present study, we aimed to derive such generalizable relationships to help predict chemical effects on ecosystems. Therefore, we conducted a meta-analysis and extracted 350 observations (effect sizes and respective chemical concentrations) from 153 published studies conducted under controlled conditions (laboratory or outdoor mesocosms) that cover a wide range of chemical use groups (metals, pesticides, and pharmaceuticals) and five ecosystem functions (community respiration, organic matter decomposition, nutrient cycling, photosynthesis, and primary production). In addition, we contextualized the observed effects within an environmental risk assessment framework by calculating risk quotients (RQs), relating the observed effects to regulatory acceptable concentrations (RACs) provided by the German Federal Environment Agency. We hypothesized that where chemicals target similar organism groups, generalizable relationships between the chemical stress intensity (standardized by their toxic potency) and rate of ecosystem function can be established (H1). Furthermore, we hypothesized that the relationship between chemical stress intensity and the ecosystem function organic matter decomposition by microbial decomposers would be less pronounced than for other functions studied (H2). This is because toxicity data for microbial decomposers (bacteria and fungi) are largely lacking, hampering a reliable characterisation of toxic potency towards them.

## 2. Materials and methods

### 2.1 Literature collection and inclusion criteria

We searched across Web of Science and Scopus to compile literature concerning the effects of chemical use groups (pesticides, pharmaceuticals, and metals) on five ecosystem functions (community respiration, nutrient cycling, organic matter decomposition, photosynthesis, and primary production). The search was limited to literature published between January 2000 and December 2024 to primarily focus on current-use pesticides and pharmaceuticals as well as metals. These chemical use groups were pre-selected because an initial screening showed that data would be too scarce for a meta-analysis of other chemical types, such as biocides. Our search string was constructed using a combination of freshwater habitat-related terms (freshwater* OR stream* OR river* OR pond* OR lake*), chemical stressor-related terms (chemical stress* OR contaminant* OR pollutant* OR toxicant* OR pesticide* OR heavy metal* OR metal* OR fungicide* OR herbicide* OR insecticide* OR pharmaceut*), and ecosystem function-related terms (ecosystem function* OR photosynth* OR primary product* OR nutrient cycl* OR organic matter decomp* OR leaf breakdown OR decomposition* OR respirat*). The asterisks (*) were employed as wildcards, capturing all terms sharing the same root. Moreover, we conducted footnote chasing by screening the reference lists of relevant articles for further literature within the considered time period. The selection process for studies encompassed three stages. Initially, titles were screened to identify suitable studies. Subsequently, the abstracts were evaluated, leading to the exclusion of non-matching studies. Lastly, the full texts of selected studies were assessed.

Studies were included based on a set of criteria: (1) laboratory-based studies and outdoor mesocosm studies; we deliberately excluded field studies from further analyses, as these represented a minor share of our dataset (*n* = 60 across all ecosystem functions), of which a major fraction (>90%) did not meet one or multiple of the following inclusion criteria, (2) focus on species, populations, or communities dwelling within freshwater habitats (excluding groundwater, marshlands, coastal waters, and marine systems), (3) experimental exposure to at least one of the pre-defined chemical types (fungicides, herbicides, insecticides, pharmaceuticals, and metals) from the list of chemical use groups above (excluding eutrophication, salinization, and acidification), (4) presence of a control treatment (absence of chemical stress), and (5) measurement of the designated ecosystem functions as well as uncertainty around the central tendency of the measurement. Furthermore, we only included studies that addressed direct effects on the organisms under scrutiny, thereby excluding indirect effects (e.g. shifts in the palatability of leaf material for detritivores due to direct effects on leaf-associated microorganisms). We excluded studies on teleost fish, as they do not contribute directly to the ecosystem functions examined here. An overview of the selected studies and moderator variables (see below) will be provided on Zenodo upon article acceptance.

### 2.2 Data extraction and effect size calculation

We extracted central tendencies, uncertainty estimators, sample sizes, chemical concentrations, and moderator variables (see below) from the text, tables, and figures of the selected studies. For cases where the data were presented in figures, we utilized the “digitize” package (Poisot, 2011) within the open-source software R (version 4.4.0 for Mac OS X; R Core Team, R Core Team, 2024).

As our response variable, we calculated the ratio of the ecosystem function observed in the stressor treatment to that in the control treatment. From each study, we extracted only the results for the concentration with the highest effect size observed per functional endpoint for the same species, population, or community. This approach was adopted to derive unequivocal cause-effect relationships and prevent pseudoreplication. Due to this method and the diverse test designs employed in individual studies, the timing of functional measurements varied.

For studies investigating community respiration, the functional indicators included measurements of O_2_-consumption (e.g., mg O_2_ g^-1^ dry weight h^-1^) and CO_2_-production (e.g., µg CO_2_ g^-1^ dry weight h^-1^). In the context of organic matter decomposition, the indicators consisted of breakdown rates (*k* (d^-1^)) and mass loss (e.g., mg d^-1^) of leaf material, mediated by microbial decomposers or a combination of microbial decomposers and detritivorous macroinvertebrates (decomposer-detritivore-systems) as endpoints. As the literature search on nutrient cycling only resulted in data for nitrogen cycling, the indicators encompassed anaerobic NH ^+^ oxidation rates (g-N g-volatile suspended solids^-1^ d^-1^), NO - and NO - turnover rates (e.g., mg NO_2_-N mg^-1^ d^-1^), and N_2_-production (e.g., N_2_ g^-1^ h^-1^). In the case of photosynthesis, we determined effect sizes based on maximum and effective photosystem II quantum yields (F_v_/F_m_ and *Φ*PSII, respectively) and O_2_-production (e.g., mg O_2_ g^-1^ dry weight h^-1^). Finally, for primary production, the effect sizes related to fixed carbon quantities (mg ^14^C g^-1^ C h^-1^) and changes in the standing biomass (e.g., g C h^-1^). Studies that exclusively reported dissolved O_2_ as a response variable were excluded, as net dissolved O_2_ could arise from multiple sources simultaneously, such as photosynthesis and the ambient atmosphere.

### 2.3 Calculation of toxic units and risk quotients

The toxic effects caused by different chemical stressors vary for a given test concentration, due to differing ecotoxicological potentials, making direct comparisons challenging. Ideally, we would have used the HC_50_ – the hazardous concentration of a stressor at which 50% of the species are exposed to a concentration above their EC_50_ (the concentration at which an effect is observed in 50% of exposed organisms) derived from a species sensitivity distribution (SSD) – as a benchmark for the toxicity of individual stressors.

However, the calculation of SSDs is not reliable if less than 10 data points are available (Hiki and Iwasaki, 2020), which was the case for >90% (especially for macrophytes) of substances. Therefore, we normalized the test concentrations of individual chemical stressors to their acute effects by calculating toxic units (TUs). Ideally, TUs would be computed for each combination of chemical stressor and organism included in the selected literature. However, given substantial data gaps, we relied on surrogate species results to facilitate comparisons between different stressors. Unlike previous studies that sought to establish generalized chemical stress–ecosystem function relationships (e.g., Peters et al., 2013), we avoided confining the surrogate species selection to conventional test species as stipulated by chemical regulations (i.e., *Daphnia magna*, *Lemna* sp., and *Raphidocelis subcapitata*). Instead, we estimated stressor toxicity based on the EC_50_ values for the most sensitive tested freshwater algal, plant, and invertebrate species (not necessarily the conventional test species used for risk assessment) for the respective chemical stressor, because this approach was superior to using a surrogate species in a previous comparison (Schäfer et al., 2013). Notwithstanding, this approach may incur some variance given its dependency on the testing effort for each chemical and the selected test species available in the literature. We employed these EC_50_ values to compute toxic units (TU) defined here as the logarithmic ratio between experimental exposure concentrations and a defined effective concentration:

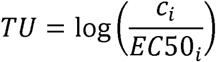

where *c* represents the concentration of a toxicant *i* and EC50*_i_* is the median effective concentration of the respective toxicant for the most sensitive species. In case of chemical mixtures (23% of cases), we employed the EC_50_ values to compute sum toxic units (sumTU) defined as the logarithmic sum of the ratios between exposure concentrations and a defined effective concentration:

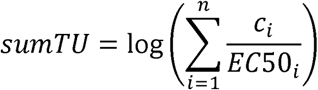

where *n* gives the number of toxicants that caused an effect in the respective ecosystem function. Although the assumption of concentration addition underlying the sumTU concept cannot be guaranteed – given that mixture components often have distinct MoAs – previous research suggests that the sumTU can reliably represent mixture toxicity, even when MoAs differ (cf. Escher et al., 2020). EC_50_ values were extracted from various sources including Standartox (Scharmüller et al., 2020), the Pesticide Property Database (Lewis et al., 2016), the Veterinary Substance Database (Lewis et al., 2011), the Norman Database (https://www.norman-network.com/nds/), and WikiPharma (Molander et al., 2009), with most of the EC_50_ being related to mortality (invertebrates) or growth (algae, macrophytes). We primarily extracted EC_50_ values corresponding to a 48-hour exposure duration or, when not available, the closest exposure time (algae: 15-120 h, invertebrates: 48-96 h, macrophytes: 15-120 h).

In addition, we extracted RACs for pesticides – used as benchmarks in environmental risk assessment – from the Information System Ecotoxicology and Environmental Quality Targets (ETOX), provided by the German Federal Environment Agency. We selected German data for its comprehensiveness and to avoid mixing data from different agencies, which may derive RACs with slightly different approaches, thereby incurring variability. RQs were calculated as the ratio between exposure concentrations and the respective RAC:

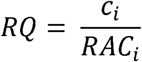

where *c* represents the concentration of a toxicant *i* and RAC*_i_* is the corresponding regulatory acceptable concentration. In case of toxicant mixtures, we employed the combined risk quotient that sums up the RQs of individual pesticides, assuming that the effects of mixtures are additive. As such benchmarks are lacking for pharmaceuticals and metals, RQ calculations could not be performed for these chemical use groups. A RQ > 1 means that the exposure exceeded the RAC. All EC_50_ values, the respective species, RACs, measured endpoints and their sources will be provided on Zenodo upon article acceptance.

### 2.4 Moderator variables

Besides central tendencies, uncertainty estimators, sample sizes, and chemical concentrations, we extracted seven moderator variables: (1) chemical type with levels: (a) fungicide, (b) herbicide, (c) insecticide, (d) pesticide mixture, (e) pharmaceutical, (f) metal, (g) metal mixture, and (h) other (such as biocides); (2) classification of the MoAs extracted from Lewis et al. (2016) or Molander et al. (2009); (3) organism group with levels: (a) algae (including cyanobacteria and phytoplankton), (b) macrophytes, (c) microbial decomposers (bacteria and fungi), (d) joint systems of microbial decomposers and invertebrate decomposers, hereafter termed decomposer-detritivore-systems; (4) exposure time as a continuous variable in days; (5) pH as a continuous variable; (6) test temperature as a continuous variable in °C; (7) study system with levels: (a) laboratory and (b) mesocosm.

### 2.5 Data analysis

To identify the moderator variables affecting the assessed ecosystem functions and establish relationships between chemical stressors and ecosystem functions, we performed separate data analyses for each of the included ecosystem functions. We used the calculated effect sizes (along with the respective standard errors of the effects) as the response variables, while including the sumTUs and moderator variables as predictors. Initially, we checked for multi-collinearity by examining correlations (r ≥ 0.8) among our predictors, ensuring that the meta-regression coefficient estimates are robust. Subsequently, we identified the best-fitting models using the “multimodel.inference” function in the R package “dmetar” (Harrer et al., 2019) based on corrected Akaike Information Criteria. Using the results of the multimodel inference, we build models to evaluate the individual impact of the identified predictors and their interactions. We achieved this by fitting meta-regression models using the R package “metafor” (Viechtbauer, 2010). We employed the “maximum likelihood” estimator to assess the amount of residual heterogeneity (*τ*^2^) and applied the Knapp-Hartung method for random-effects meta-analyses (van Aert and Jackson, 2019). We conducted a permutation test running 1,000 iterations before reporting *p*-values, as recommended for meta-regression models, to control for type I error rates of frequentist meta-regressions with small sample sizes (Higgins and Thompson, 2004).

Afterwards, we fitted (quasi-)binomial Generalized Linear Models (GLMs) to explore potential dose-response relationships between ecosystem functions and sumTUs as well as RQs. Models were fitted for all stressors combined or separately for chemical types (fungicides, herbicides, insecticides, pharmaceuticals, and metals) or MoAs (only if the number of observations n ≥ 8, given the statistical limitations of small sample sizes; Jenkins and Quintana-Ascencio, 2020). To exclude the influence of extreme data points (such as very low or very high responses of ecosystem functions), we employed bootstrapping (1,000 times). Models were then run on the individual bootstrapped data sets, and we calculated the model medians to represent the central tendency of the fitted models. For statistical analyses and graphics, we used the open-source software R (R Core Team, 2024), along with the necessary add-on packages. Data and code will be freely available on Zenodo upon acceptance.

## 3. Results

### 3.1 Overview of included studies

Through our database query and footnote chasing, we identified a total of 7,661 publications, of which 7,508 publications were excluded due to violations of our predefined inclusion criteria, absence of reported effect sizes and associated uncertainties, or the inability to calculate sumTUs due to a lack of toxicity data. Ultimately, we focused on 153 studies that met our criteria, from which we extracted 350 effect sizes (# of observations), uncertainty estimators, sumTUs, and moderator variables (see Fig. A1 for details). Among the selected studies, 37% of observations each focused on organic matter decomposition and photosynthesis, while 21% addressed primary production. Due to limited representation (5%), community respiration and nutrient cycling were excluded from further analysis. Pesticides were the most frequently studied chemical stressors (66%, n=232), comprising fungicides (n=67), herbicides (n=86), insecticides (n=43), and pesticide mixtures (n=36). Metals and their mixtures accounted for 26% (n=91), and pharmaceuticals for 8% (n=27; Fig. 1). SumTU values ranged from –5.13 to 2.68 (metals), –7.60 to 3.99 (pesticides), and –5.68 to 5.59 (pharmaceuticals), with 0 indicating toxicity equivalent to the EC50 of the most sensitive species (Table 1).

**Figure 1:**
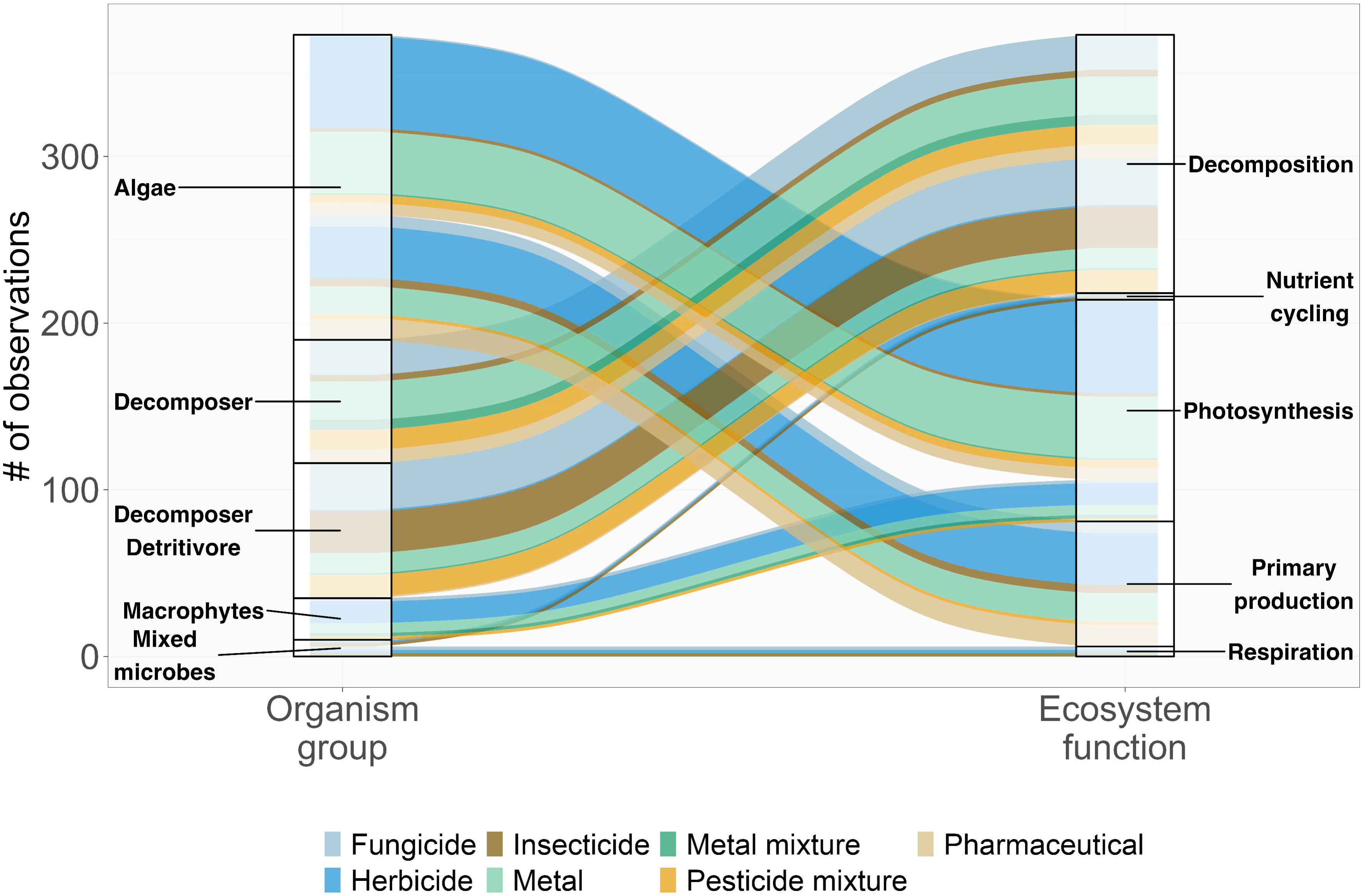
Alluvial plot displaying the extracted effect data. The size of each box is proportional to the number of extracted observations per organism group (left) or ecosystem function (right). The link width is proportionate to the number of observations per chemical type (indicated by color).

**Table 1:** Descriptive statistics on sumTU values for the individual chemical types (*n* = 350 effects sizes or # of observations).

| Chemical type | $n$ | Mean | Min | 1. Quartile | 2. Quartile<br>(Median) | 3. Quartile | 4. Quartile |
| --- | --- | --- | --- | --- | --- | --- | --- |
| Fungicides | 67 | -0.01 | -7.60 | -0.82 | 0.11 | 0.59 | 2.31 |
| Herbicides | 86 | 1.26 | -3.03 | 0.87 | 1.39 | 1.57 | 3.99 |
| Insecticides | 43 | 0.34 | -4.93 | -0.90 | 0.75 | 2 | 3.60 |
| Pesticide mixtures | 36 | 0.09 | -2.31 | -1.00 | 0.08 | 0.94 | 2.78 |
| Pharmaceuticals | 27 | 0.08 | -5.68 | -1.05 | 0.26 | 0.87 | 5.59 |
| Metals | 81 | 0.33 | -5.13 | -0.49 | 0.56 | 1.66 | 2.68 |
| Metal mixtures | 10 | 0.86 | -1.26 | 0.55 | 0.99 | 1.43 | 1.77 |

### 3.2 Influence of moderator variables and relationships between sumTU and effects on ecosystem functions

#### 3.2.1 Organic matter decomposition

For the ecosystem function organic matter decomposition, our best-fit model included two significant predictors: “organism group” (*p* = 0.001), which categorized microbial decomposers alone and decomposer-detritivore-systems, and “pH” (*p* = 0.007). Additionally, “organism group” and “sumTU” exhibited a significant interaction (*p* = 0.039). This model accounted for a substantial portion of the variance (meta-regression: R^2^ = 58.93%, τ^2^ = 0.0434; Table 2). We observed a general dose-response relationship for organic matter decomposition across all groups of chemicals for studies examining decomposer-detritivore systems (as indicated by the GLM for sumTU and the respective ecosystem function; *p* = 0.022; *n* = 93, Fig. 2a). However, when focusing on microbial decomposers alone, we did not find a consistent relationship between sumTU and organic matter decomposition (*p* = 0.801; *n* = 82; Fig. 2b), nor did we find such a relationship for any individual chemical type or MoA (*p* ≥ 0.288; *n* = 8-22).

**Figure 2:**
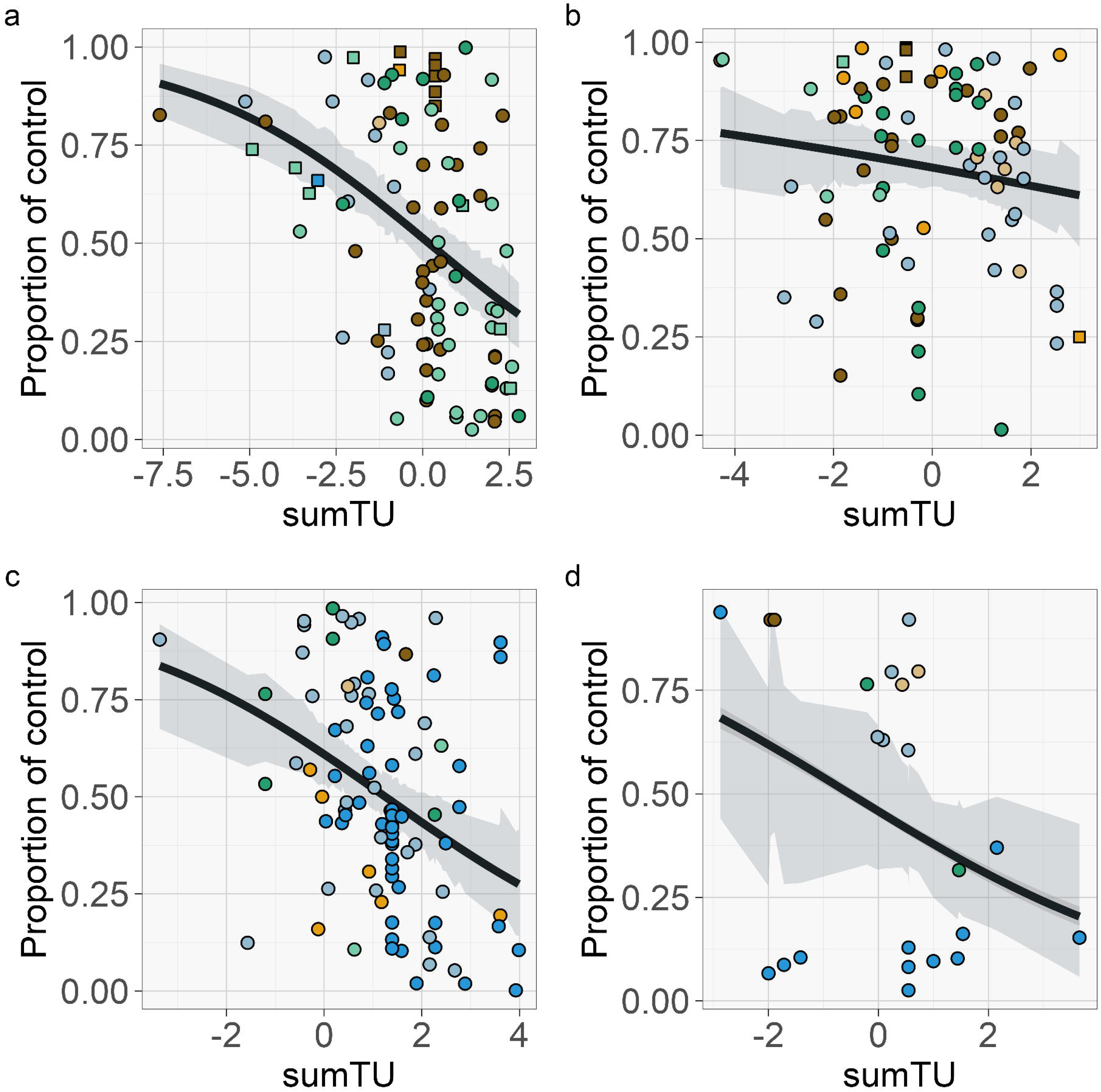
Relationships between sumTU and (a) organic matter decomposition by decomposer-detritivore-systems, (b) organic matter decomposition by microbial decomposers, (c) photosynthesis by algae, and (d) photosynthesis by macrophytes, shown as the proportion of the functioning in the stressor-treatment compared to the stressor-free control. Colors code chemical types (see Fig. 1), and shapes represent the study system, namely laboratory (circles) and micro-/mesocosms (squares). The shaded area represents the 95% confidence interval based on the bootstrapped data, while the black line represents the median model. Please note the differences in the x-axis scale.

**Table 2:**
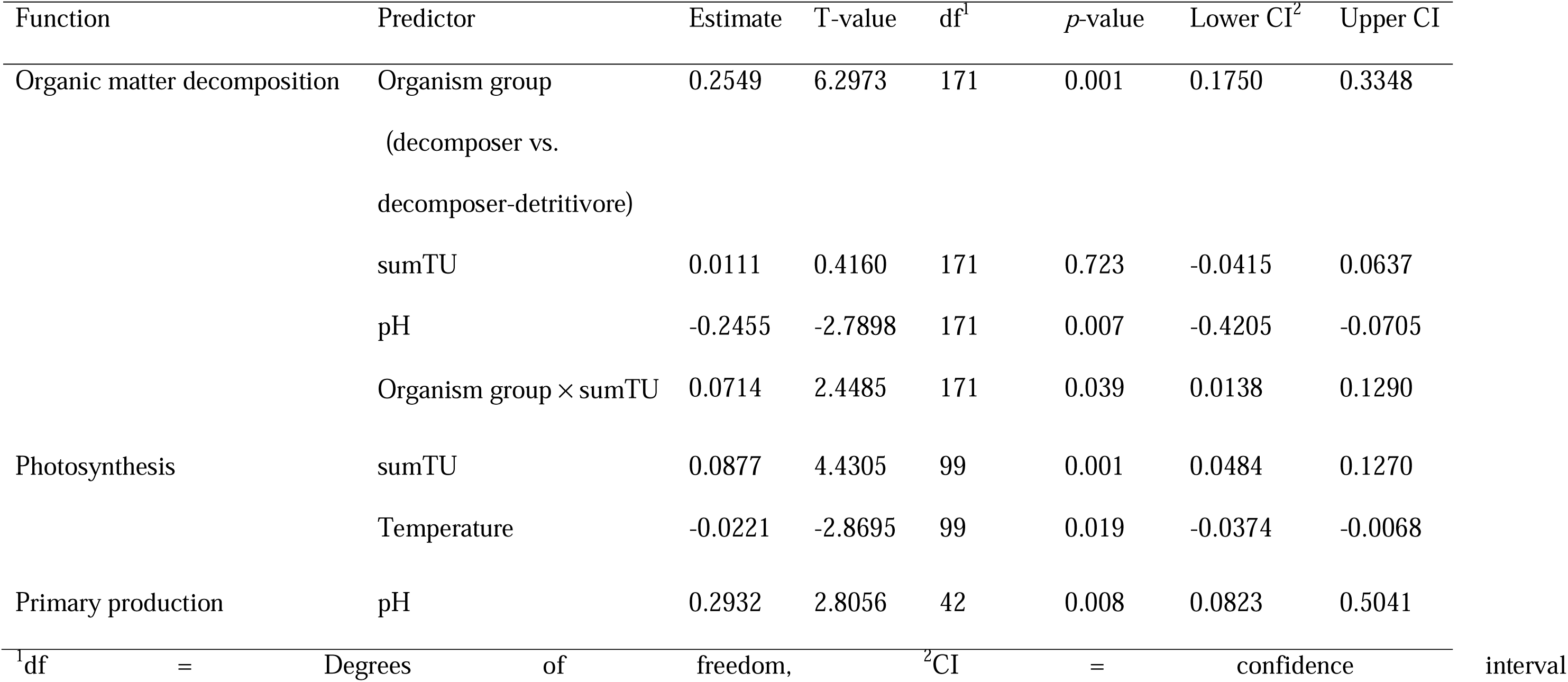
Outcome of meta-regression models assessing the influence of moderator variables on ecosystem functions.

#### 3.2.2 Photosynthesis

For the ecosystem function photosynthesis, our best-fit model included two significant predictors: “sumTU” (*p* = 0.001) and “temperature” (*p* = 0.023), collectively explaining a portion of the variance (R^2^ = 18.22%, τ^2^ = 0.0707; Table 2). In contrast to organic matter decomposition, we did not observe an influence of the parameter “organism group” (distinguishing between algae and macrophytes; Fig. 2c&d). When we examined the effects of individual chemical types or MoA on specific organism groups, no clear dose-response relationships were evident (*p* ≥ 0.210; *n* = 12-50).

#### 3.2.3 Primary production

For the ecosystem function primary production, our best-fit model included “pH” (*p* = 0.008) as significant predictor, explaining a portion of the variance (R^2^ = 16.06%, τ^2^ = 0.0437; Table 2), but not “sumTU”. When we examined the effects of individual chemical types or MoA on algae (lack of data on macrophytes), no clear dose-response relationships were evident (*p* ≥ 0.285; *n* = 12-24).

### 3.3 Relationships between risk quotients for regulatory thresholds and effects on ecosystem functions

We did not find a general relationship between RQs and ecosystem functioning related to organic matter decomposition across all chemical groups, neither when focusing on decomposer–detritivore systems (GLM of logCC(RQ) and the respective ecosystem function; p = 0.933; n = 71) nor when focusing on decomposers only (p = 0.335; n = 41). For photosynthesis, a general relationship was found for algae (p = 0.008; n = 35), whereas no such relationship was observed for studies examining primary production (p = 0.989; n = 15; Fig. 3).

**Figure 3:**
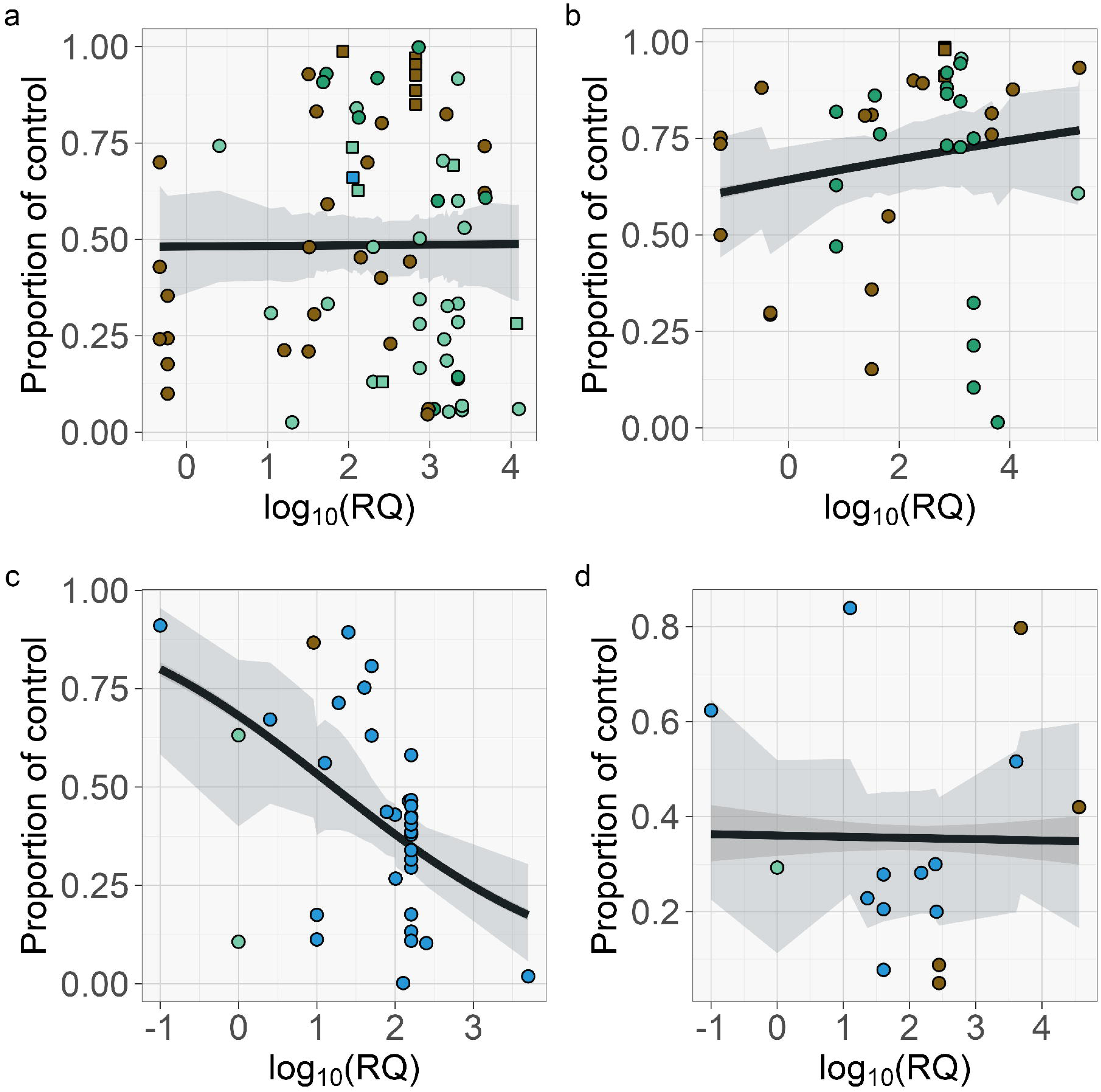
Relationships between logCC(RQ) and (a) organic matter decomposition by decomposer-detritivore-systems, (b) organic matter decomposition by microbial decomposers, (c) photosynthesis by algae, and (d) photosynthesis by macrophytes, shown as the proportion of the functioning in the stressor-treatment compared to the stressor-free control. Colors code chemical types (see Fig. 1), and shapes represent the study system, namely laboratory (circles) and micro-/mesocosms (squares). The shaded area represents the 95% confidence interval based on the bootstrapped data, while the black line represents the median model. Please note the differences in the x-axis scale.

## 4. Discussion

### 4.1 Influence of moderator variables and relationships between sumTU and effects on ecosystem functions

To the best of our knowledge, our study is among the first in establishing relationships between chemical stressors and ecosystem functions. This represents a crucial stride, as such relationships can ultimately aid in predicting the toxicity of chemicals that have not yet been thoroughly studied, although some degree of uncertainty remains due to the broad range of MoAs. Nevertheless, the transferability and predictive capacity of these relationships could potentially reduce costs and efforts, diminish the reliance on animal testing, and mitigate the suffering of living organisms – an issue that has received considerable attention in the last decade (Burden et al., 2015).

In accordance with our first hypothesis (H1), we found a few generalizable relationships between the toxic potency of chemicals that target similar organism groups and the rate of ecosystem functions. These relationships exhibited an inverted S-shaped monotonicity, signifying that the measured parameters declined as the toxic potency of the chemical stressors increased. Our findings therefore align with the dose-response patterns frequently observed in the field of ecotoxicology (Newman, 2009). The S-shape observed in these relationships suggests that organisms carry specific defense or repair mechanisms that provide protection against a certain amount of a chemical stressor, up to a critical threshold level. Beyond this threshold, organisms become incapable of defending or repairing themselves effectively. Consequently, the biological response to chemical stressors showed a graded response due to the evolutionary adaptive capabilities of organisms to react to a range of low doses. The absence of relationships between toxic potency and ecosystem functioning apart from those identified for organic matter decomposition and photosynthesis may be explained by three factors. First, the toxic pressure from certain chemical groups may have been below thresholds triggering adverse effects. Second, bias related to our toxicity estimation particularly for microbial decomposers may have obscured relationships (H2). Third, given considerable variation in the obtained regression relationships, incurred by pooling studies with very different conditions and chemicals with differing MoAs, may have hidden a potential relationship.

For the ecosystem function organic matter decomposition by decomposer-detritivore-systems, we observed a 50% reduction occurring at a sumTU of approximately 0. This finding aligns with a field study that observed a substantial decrease in organic matter decomposition by decomposer-detritivore-systems of approximately 50% at a an even lower log(TU_D,magna_) of 0.01 (Schäfer et al., 2012a). Consequently, there is a good overlap of field data and the relationships between chemical exposures and their effect on organic matter decomposition observed in our study, despite the inherent challenges of comparing data sets – for instance studies mainly focused on individual chemicals in laboratory or mesocosm settings in comparison to complex mixtures in the field – highlighting the value of our results.

Despite this good overlap of field data and the chemical stress-ecosystem function relationship derived here, it is essential to acknowledge that the heavy emphasis on specific chemical use groups in research often hampers the broader applicability of laboratory findings to real-world conditions. Over the past 25 years, a predominant focus has been on metals, pesticides (particularly herbicides), and pharmaceuticals. However, these chemical use groups constitute only a small fraction of the vast array of chemicals present in the environment, given that approximately 350,000 chemicals are registered for production and use (Wang et al., 2020). Studies employing bioassays have revealed that target chemicals can only account for a small fraction of the overall response to chemicals (as low as < 0.1 %; Escher et al., 2013), suggesting that numerous ecotoxicologically important compounds exist within freshwater ecosystems (Escher et al., 2013; Neale et al., 2020). In addition to the emphasis on pesticides, metals, and pharmaceuticals, only around one quarter of the included studies have considered that chemicals often co-occur as mixtures. Typically, the effect of chemicals in mixtures is additive, but in a relevant proportion of cases, the simultaneous presence of multiple chemicals can result in stronger (synergistic) or weaker (antagonistic) effects compared to the assumption of additive effects (cf. Warne and Hawker, 1995). A recent review, encompassing various chemical groups, indicated an average of 20% of cases with synergism and 15% of cases with antagonism (Martin et al., 2021). Another review reported synergistic effects in 3% to 26% of cases involving binary mixtures, with the prevalence varying by chemical group (Cedergreen, 2014). Furthermore, freshwater ecosystems are typically subject to multiple stressors concurrently (Birk et al., 2020), which can lead to interactions between chemical and non-chemical stressors (Liess et al., 2016). To provide a more comprehensive understanding of chemical effects in freshwater ecosystems and derive additional chemical stress-ecosystem function relationships, it is essential to broaden the scope. This entails identifying ecotoxicologically relevant chemical groups, assessing interactions within chemical mixtures, and considering the context of multiple (non-chemical) stressors in the ecosystem.

Furthermore, finding chemical stressor-ecosystem function relationships can be confounded by abiotic factors, such as pH (Ragnarsdottir, 2000; Wang et al., 2016) and temperature (Soares et al., 2020), which influence the fate and toxicity of metals and organic chemicals in aquatic environments. This phenomenon is often referred to as “context dependency” (Catford et al., 2022). While pH and temperature had direct effects on ecosystem functions, which are discussed below, no interactions between pH or temperature and other moderator variables such as the toxic potency were observed. However, such interactions have been observed earlier. For instance, a meta-analysis revealed a substantial influence of pH on metal toxicity (Wang et al., 2016). This can be explained by the pH-dependency of metal speciation, which in turn affects metal bioavailability. Furthermore, pesticides can be ionizable compounds (Ragnarsdottir, 2000), and their degradation and subsequent toxicity are influenced by pH as well as molecular structural properties (Fenner et al., 2013). Temperature also influences the effects of chemical stressors through two key concepts. The “climate-induced toxicant sensitivity” (CITS) concept describes how climate-related environmental factors alter chemical toxicity, while the “toxicant-induced climate change sensitivity” (TICS) concept explains how pesticide exposure increases susceptibility to climate change (Verheyen et al., 2022). These concepts are interconnected, as global warming may enhance pesticide toxicity (CITS), which in turn could further reduce heat tolerance (TICS; Verheyen et al., 2022). Moreover, aside from their role in modifying chemical bioavailability, speciation, and concentration, abiotic conditions have been shown to increase species sensitivity by up to 30-fold in certain experiments (Liess et al., 2001; Liess and Beketov, 2011).

Additionally, biotic factors can impede the derivation of chemical stressor-ecosystem function relationships. However, the relevance of these factors largely depends on the ecosystem function under investigation and the design of the experimental approach. Laboratory experiments typically involve the use of species that are maintained in culture over extended periods, often under conditions that differ from their natural habitats (e.g., variations in light, temperature, and pH) and that may lay outside their niche width and optima. Thus, these conditions affect the survival, growth, reproduction, and functioning of species, and thereby the related ecosystem functions (Blackman, 1905). Indeed, pH and temperature significantly influenced organic matter decomposition, photosynthesis, and primary production. These findings suggest a mismatch between species-specific requirements and the laboratory conditions under which the respective experiments were conducted. In studies that make use of complex natural communities rather than single species tests (such as microbially-mediated organic matter decomposition), additional biotic factors can introduce uncertainties into the derivation of chemical stressor-ecosystem function relationships. These studies typically involve communities with different compositions, and these structural differences have been demonstrated to influence the sensitivity of communities to chemical stress (Beketov et al., 2008; Feckler et al., 2018), for instance due to functional similarity among species that has not been considered in the present study (Louca et al., 2018). Consequently, these variations can lead to differing effects on ecosystem functions due to differences in the capacity of communities to compensate for species loss caused by chemical stressors (Cadotte et al., 2011).

### 4.2 Study bias and confounding factors in the present meta-analysis

We specifically chose biologically-mediated ecosystem functions for our analysis, focusing on those we deemed most ecologically relevant within freshwater ecosystems.

Among the five selected ecosystem functions, primary production and photosynthesis by algae and macrophytes were notably prevalent. This prevalence can be attributed to several factors. First, these functions are likely to be highly responsive to the most studied group of pesticides, namely herbicides (Hedlund et al., 2020), as well as pharmaceuticals. Cyanobacteria, as photosynthetic prokaryotes, are considered sensitive to antibiotics due to their close relationship with pathogenic bacteria (González-Pleiter et al., 2013), and green algae and macrophytes can also be affected by antibiotics because of the prokaryotic origin of their semi-autonomous cell organelles (Nie et al., 2009). Second, maintaining algal cultures and macrophytes in laboratory settings is relatively straightforward, and measurement techniques for primary production and photosynthesis enable rapid quantitative assessment.

Despite their importance for freshwater ecosystem functioning, their potential of being used as indicators of streams’ ecological status and in the assessment of streams’ restoration (Klocker et al., 2009; Udy et al., 2006; Young et al., 2008), only a minor share of studies addressed chemical stressor effects on microbial community respiration and nutrient cycling. Unlike for algae, standardized protocols for culturing heterotrophic microbial communities (bacteria and fungi) in the laboratory are largely lacking, hindering their use in research and complicating comparisons between studies. Moreover, studies assessing ecosystem functions through microbial decomposers (approximately 24 ± 2 days; mean ± standard error) required considerably longer exposure times compared to those focusing on primary production and photosynthesis by algae (approximately 5 ± 0.5 days), increasing the labor-intensity. Nevertheless, protocols to quantify various processes related to community respiration and nutrient cycling have been developed and are frequently applied in field studies to assess stream ecosystem health (von Schiller et al., 2017). While these protocols are similarly complex and costly as other methods for quantifying ecosystem functions, they could be partially automated for use in laboratory studies (von Schiller et al., 2017). Thus, we recommend their application in future laboratory studies to broaden our mechanistic understanding of how chemical stressors impact ecosystem functions.

Lastly, the taxonomic mismatch between the most sensitive tested species used for calculating TUs and the species assessed for the chemical stress–ecosystem function relationship likely introduced some uncertainty. However, we deliberately used the most sensitive reported species for each chemical stressor to provide a conservative estimate of their effects on ecosystem functions. Given that these sensitive species are present in freshwater communities and are affected by chemical exposure, the observed effects are expected to be comparable to those identified in this meta-analysis (cf. Schäfer et al., 2012b).

### 4.3 Relationships between risk quotients for regulatory thresholds and effects on ecosystem functions

Only one of four relationships between RQs of RACs and effects on ecosystem functions was significant. Ideally, ecosystem functions would be at control levels for RQs < 1 and only decline when this threshold is exceeded, equaling the exceedance of RACs. The fact that ecosystem functions were frequently much lower than control levels for RQs < 1 and that only one consistent relationship could be established questions the protectiveness of RACs for ecosystem functions and their accuracy in reflecting real-world ecological dynamics. Several factors likely explain this shortcoming of RACs. First, RACs are derived from toxicity data that relate to single-species laboratory or multispecies mesocosm studies. The relationship between single-species experiments and ecosystem functions can vary strongly because of variability in the contribution of the single species to the function under scrutiny and complex interspecies interactions not accounted for (Kraft et al., 2015). Likewise, the responses to chemicals can vary strongly between mesocosm studies due to experimental conditions (e.g., Beketov and Lies, 2008) and the most sensitive response variable selected for RAC derivation may vary regarding its relationship with ecosystem functioning. Second, the thresholds derived from current risk assessments for single species or populations may underestimate functional impairments originating from sublethal effects. Consequently, key ecosystem processes may be disrupted even when stressor concentrations remain below levels deemed “acceptable” for non-functional responses. Third, the relationship between risk quotients for RAC and ecosystem functioning may be nonlinear and moderated by non-additive mixture effects that are not adequately captured by current risk assessment approaches.

## 5. Conclusions

Along with the ever-increasing production values and usage of new chemicals, it becomes insurmountable to experimentally study the effects of all chemicals alone and in combination with other chemicals and stressors in a wide range of ecosystems, making generalizations invaluable. In this context, our analyses revealed the potential of deriving chemical stress-ecosystem function relationships using currently available data that show adverse effects for certain ecosystem functions. Nevertheless, we identified several important research gaps – partially beyond the scope of the present study – that should be considered in the future to support deriving more specific (e.g., in terms of MoA) chemical stress-ecosystem function relationships and ultimately facilitate the prediction of chemical effects. As we showed in our analysis, chemicals of certain groups as well as specific ecosystem functions were underrepresented in the extracted literature published between 2000 and 2024. Therefore, data likely remains insufficient for important but understudied chemical groups and ecosystem functions, hampering a more complete evaluation of chemical stress. Furthermore, we showed that the context-dependency of chemical stress effects on ecosystem functions is largely ignored. Therefore, future laboratory research should proactively consider a series of abiotic and biotic factors as well as the composition of stressors (single stressors vs. diverse chemical mixtures; combination of chemical and non-chemical stressors) to ensure the transferability of the data to various field situations. Ideally, such studies should simulate different climatic and biogeographical regions as well as chemical exposure histories as these factors were shown to determine the stress response (Zubrod et al., 2019; Sinclair et al., 2024). In addition, laboratory studies should be appended by semi-field and field studies as verification and validation benchmarks to assess the accuracy and reliability of laboratory data for field situations. Finally, there is a disconnect between regulatory ecological quality targets and ecological outcomes, highlighting a need to re-evaluate risk assessment approaches if they are supposed to be ecologically meaningful and protective of ecosystem functions.

## Data accessibility statement

All data used for the analyses of the present study were extracted from previously published articles. All data and R code used in this paper will be uploaded to Zenodo upon article acceptance.

## References

Balian, E.V., Segers, H., Martens, K., Lévéque, C., 2008. The Freshwater Animal Diversity Assessment: an overview of the results, in: Balian, E.V., Lévêque, C., Segers, H., Martens, K. (Eds.), Freshwater Animal Diversity Assessment, Developments in Hydrobiology. Springer Netherlands, Dordrecht, pp. 627–637. 10.1007/978-1-4020-8259-7_61

Battin, T.J., Lauerwald, R., Bernhardt, E.S., Bertuzzo, E., Gener, L.G., Hall, R.O., Hotchkiss, E.R., Maavara, T., Pavelsky, T.M., Ran, L., Raymond, P., Rosentreter, J.A., Regnier, P., 2023. River ecosystem metabolism and carbon biogeochemistry in a changing world. Nature 613, 449–459. 10.1038/s41586-022-05500-8

Beaumelle, L., De Laender, F., Eisenhauer, N., 2020. Biodiversity mediates the effects of stressors but not nutrients on litter decomposition. eLife 9, e55659. 10.7554/eLife.55659

Beketov, M.A., Liess, M., 2008. Variability of pesticide exposure in a stream mesocosm system: Macrophyte-dominated vs. non-vegetated sections. Environmental Pollution 156, 1364–1367. 10.1016/j.envpol.2008.08.014

Beketov, M.A., Schäfer, R.B., Marwitz, A., Paschke, A., Liess, M., 2008. Long-term stream invertebrate community alterations induced by the insecticide thiacloprid: Effect concentrations and recovery dynamics. Sci. Total Environ. 405, 96–108. 10.1016/j.scitotenv.2008.07.001

Bernhardt, E.S., Rosi, E.J., Gessner, M.O., 2017. Synthetic chemicals as agents of global change. Front. Ecol. Environ. 15, 84–90. 10.1002/fee.1450

Birk, S., Chapman, D., Carvalho, L., Spears, B.M., Andersen, H.E., Argillier, C., Auer, S., Baattrup-Pedersen, A., Banin, L., Beklioğlu, M., Bondar-Kunze, E., Borja, A., Branco, P., Bucak, T., Buijse, A.D., Cardoso, A.C., Couture, R.-M., Cremona, F., de Zwart, D., Feld, C.K., Ferreira, M.T., Feuchtmayr, H., Gessner, M.O., Gieswein, A., Globevnik, L., Graeber, D., Graf, W., Gutiérrez-Cánovas, C., Hanganu, J., Işkın, U., Järvinen, M., Jeppesen, E., Kotamäki, N., Kuijper, M., Lemm, J.U., Lu, S., Solheim, A.L., Mischke, U., Moe, S.J., Nõges, P., Nõges, T., Ormerod, S.J., Panagopoulos, Y., Phillips, G., Posthuma, L., Pouso, S., Prudhomme, C., Rankinen, K., Rasmussen, J.J., Richardson, J., Sagouis, A., Santos, J.M., Schäfer, R.B., Schinegger, R., Schmutz, S., Schneider, S.C., Schülting, L., Segurado, P., Stefanidis, K., Sures, B., Thackeray, S.J., Turunen, J., Uyarra, M.C., Venohr, M., von der Ohe, P.C., Willby, N., Hering, D., 2020. Impacts of multiple stressors on freshwater biota across spatial scales and ecosystems. Nat. Ecol. Evol. 4, 1060–1068. 10.1038/s41559-020-1216-4

Blackman, F.F., 1905. Optima and limiting factors. Ann. Bot. 19, 281–295.

Burden, N., Sewell, F., Chapman, K., 2015. Testing chemical safety: what is needed to ensure the widespread application of non-animal approaches? PLOS Biology 13, e1002156. 10.1371/journal.pbio.1002156

Cadotte, M.W., Carscadden, K., Mirotchnick, N., 2011. Beyond species: functional diversity and the maintenance of ecological processes and services. J. Appl. Ecol. 48, 1079– 1087. 10.1111/j.1365-2664.2011.02048.x

Cardinale, B.J., Duffy, J.E., Gonzalez, A., Hooper, D.U., Perrings, C., Venail, P., Narwani, A., Mace, G.M., Tilman, D., Wardle, D.A., Kinzig, A.P., Daily, G.C., Loreau, M., Grace, J.B., Larigauderie, A., Srivastava, D.S., Naeem, S., 2012. Biodiversity loss and its impact on humanity. Nature 486, 59–67. 10.1038/nature11148

Catford, J.A., Wilson, J.R.U., Pyšek, P., Hulme, P.E., Duncan, R.P., 2022. Addressing context dependence in ecology. Trends Ecol. Evol. 37, 158–170. 10.1016/j.tree.2021.09.007

Cedergreen, N., 2014. Quantifying synergy: a systematic review of mixture toxicity studies within environmental toxicology. PLOS ONE 9, e96580. 10.1371/journal.pone.0096580

Corcoll, N., Casellas, M., Huerta, B., Guasch, H., Acuña, V., Rodríguez-Mozaz, S., Serra-Compte, A., Barceló, D., Sabater, S., 2015. Effects of flow intermittency and pharmaceutical exposure on the structure and metabolism of stream biofilms. Sci.Total Environ. 503–504, 159–170. 10.1016/j.scitotenv.2014.06.093

Crane, M., Watts, C., Boucard, T., 2006. Chronic aquatic environmental risks from exposure to human pharmaceuticals. Sci. Total Environ. 367, 23–41. 10.1016/j.scitotenv.2006.04.010

Dirzo, R., Young, H.S., Galetti, M., Ceballos, G., Isaak, N.J.B., Collen, B., 2014. Defaunation in the Anthropocene. Science 345, 401–406. https://10.1126/science.1251817

Escher, B.I., van Daele, C., Dutt, M., Tang, J.Y.M., Altenburger, R., 2013. Most oxidative stress response in water samples comes from unknown chemicals: the need for effect-based water quality trigger values. Environ. Sci. Technol. 47, 7002–7011. 10.1021/es304793h

Escher, B., Braun, G., Zarfl, C., 2020. Exploring the Concepts of Concentration Addition and Independent Action Using a Linear Low-Effect Mixture Model. Environmental Toxicology and Chemistry 39, 2552–2559. 10.1002/etc.4868

Feckler, A., Bundschuh, M., 2020. Decoupled structure and function of leaf-associated microorganisms under anthropogenic pressure: Potential hurdles for environmental monitoring. Freshw. Sci. 39, 652–664. 10.1086/709726

Feckler, A., Goedkoop, W., Konschak, M., Bundschuh, R., Kenngott, K.G.J., Schulz, R., Zubrod, J.P., Bundschuh, M., 2018. History matters: heterotrophic microbial community structure and function adapt to multiple stressors. Glob. Change Biol. 24, e402–e415. 10.1111/gcb.13859

Feckler, A., Kahlert, M., Bundschuh, M., 2015. Impacts of contaminants on the ecological role of lotic biofilms. Bull. Environ. Contam. Toxicol. 95, 421–427. 10.1007/s00128-015-1642-1

Fenner, K., Canonica, S., Wackett, L.P., Elsner, M., 2013. Evaluating pesticide degradation in the environment: blind spots and emerging opportunities. Science 341, 752–758. 10.1126/science.1236281

Ferreira, V., Koricheva, J., Duarte, S., Niyogi, D.K., Guérold, F., 2016. Effects of anthropogenic heavy metal contamination on litter decomposition in streams – A meta-analysis. Environ. Pollut. 210, 261–270. 10.1016/j.envpol.2015.12.060

González-Pleiter, M., Gonzalo, S., Rodea-Palomares, I., Leganés, F., Rosal, R., Boltes, K., Marco, E., Fernández-Piñas, F., 2013. Toxicity of five antibiotics and their mixtures towards photosynthetic aquatic organisms: Implications for environmental risk assessment. Water Res. 47, 2050–2064. 10.1016/j.watres.2013.01.020

Guo, J., Boxall, A., Selby, K., 2015. Do pharmaceuticals pose a threat to primary producers? Crit. Rev. Environ. Sci. Technol. 45, 2565–2610. 10.1080/10643389.2015.1061873

Halstead, N.T., McMahon, T.A., Johnson, S.A., Raffel, T.R., Romansic, J.M., Crumrine, P.W., Rohr, J.R., 2014. Community ecology theory predicts the effects of agrochemical mixtures on aquatic biodiversity and ecosystem properties. Ecology Letters 17, 932–941. 10.1111/ele.12295

Harrer, M., Cuijpers, P., Furukawa, T., Ebert, D.D., 2019. dmetar: Companion R Package For The Guide “Doing Meta-Analysis in R.”

Harrison, I., Abell, R., Darwall, W., Thieme, M.L., Tickner, D., Timboe, I., 2018. The freshwater biodiversity crisis. Science 362, 1369–1369. 10.1126/science.aav9242

Harrison, P.A., Berry, P.M., Simpson, G., Haslett, J.R., Blicharska, M., Bucur, M., Dunford, R., Egoh, B., Garcia-Llorente, M., Geamănă, N., Geertsema, W., Lommelen, E., Meiresonne, L., Turkelboom, F., 2014. Linkages between biodiversity attributes and ecosystem services: a systematic review. Ecosyst. Serv. 9, 191–203. 10.1016/j.ecoser.2014.05.006

Hedlund, J., Longo, S.B., York, R., 2020. Agriculture, pesticide use, and economic development: a global examination (1990–2014). Rural Sociol. 85, 519–544. 10.1111/ruso.12303

Hiki, K., Iwasaki, Y., 2020. Can We Reasonably Predict Chronic Species Sensitivity Distributions from Acute Species Sensitivity Distributions? Environ. Sci. Technol. 54, 13131–13136. 10.1021/acs.est.0c03108

Higgins, J.P.T., Thompson, S.G., 2004. Controlling the risk of spurious findings from meta-regression. Stat. Med. 23, 1663–1682. 10.1002/sim.1752

Jenkins, D.G., Quintana-Ascencio, P.F., 2020. A solution to minimum sample size for regressions. PLOS ONE 15, e0229345. 10.1371/journal.pone.0229345

Jepsen, R., He, K., Blaney, L., Swan, C., 2019. Effects of antimicrobial exposure on detrital biofilm metabolism in urban and rural stream environments. Sci. Total Environ. 666, 1151–1160. 10.1016/j.scitotenv.2019.02.254

Klocker, C.A., Kaushal, S.S., Groffman, P.M., Mayer, P.M., Morgan, R.P., 2009. Nitrogen uptake and denitrification in restored and unrestored streams in urban Maryland, USA. Aquat. Sci. 71, 411–424. 10.1007/s00027-009-0118-y

Kovalakova, P., Cizmas, L., McDonald, T.J., Marsalek, B., Feng, M., Sharma, V.K., 2020. Occurrence and toxicity of antibiotics in the aquatic environment: a review. Chemosphere 251, 126351. 10.1016/j.chemosphere.2020.126351

Kraft, N.J.B., Adler, P.B., Godoy, O., James, E.C., Fuller, S., Levine, J.M., 2015. Community assembly, coexistence and the environmental filtering metaphor. Functional Ecology 29, 592–599. 10.1111/1365-2435.12345

Kümmerer, K., 2009. Antibiotics in the aquatic environment – a review – Part I. Chemosphere 75, 417–434. 10.1016/j.chemosphere.2008.11.086

Lewis, K., Tzilivakis, J., Green, A., Warner, D.J., 2011. Veterinary Substances Database (VSDB).

Lewis, K.A., Tzilivakis, J., Warner, D.J., Green, A., 2016. An international database for pesticide risk assessments and management. Hum. Ecol. Risk Assess. 22, 1050–1064. 10.1080/10807039.2015.1133242

Liess, M., Beketov, M., 2011. Traits and stress: keys to identify community effects of low levels of toxicants in test systems. Ecotoxicology 20, 1328–1340. 10.1007/s10646-011-0689-y

Liess, M., Champeau, O., Riddle, M., Schulz, R., Duquesne, S., 2001. Combined effects of ultraviolet-B radiation and food shortage on the sensitivity of the Antarctic amphipod *Paramoera walkeri* to copper. Environ. Toxicol. Chem. 20, 2088–2092. 10.1002/etc.5620200931

Liess, M., Foit, K., Knillmann, S., Schäfer, R.B., Liess, H.-D., 2016. Predicting the synergy of multiple stress effects. Sci. Rep. 6, 32965. 10.1038/srep32965

Liess, M., Liebmann, L., Vormeier, P., Weisner, O., Altenburger, R., Borchardt, D., Brack, W., Chatzinotas, A., Escher, B., Foit, K., Gunold, R., Henz, S., Hitzfeld, K.L., Schmitt-Jansen, M., Kamjunke, N., Kaske, O., Knillmann, S., Krauss, M., Küster, E., Link, M., Lück, M., Möder, M., Müller, A., Paschke, A., Schäfer, R.B., Schneeweiss, A., Schreiner, V.C., Schulze, T., Schüürmann, G., von Tümpling, W., Weitere, M., Wogram, J., Reemtsma, T., 2021. Pesticides are the dominant stressors for vulnerable insects in lowland streams. Water Res. 201, 117262. 10.1016/j.watres.2021.117262

Louca, S., Polz, M.F., Mazel, F., Albright, M.B.N., Huber, J.A., O’Connor, M.I., Ackerman, M., Hahn, A.S., Srivastava, D.S., Crowe, S.A., Doebeli, M., Parfrey, L.W., 2018. Function and functional redundancy in microbial systems. Nature Ecol. Evol. 2, 936–943. 10.1038/s41559-018-0519-1

Malaj, E., Ohe, P.C. von der, Grote, M., Kühne, R., Mondy, C.P., Usseglio-Polatera, P., Brack, W., Schäfer, R.B., 2014. Organic chemicals jeopardize the health of freshwater ecosystems on the continental scale. PNAS 111, 9549–9554. 10.1073/pnas.1321082111

Martin, O., Scholze, M., Ermler, S., McPhie, J., Bopp, S.K., Kienzler, A., Parissis, N., Kortenkamp, A., 2021. Ten years of research on synergisms and antagonisms in chemical mixtures: a systematic review and quantitative reappraisal of mixture studies. Environ. Int. 146, 106206. 10.1016/j.envint.2020.106206

Molander, L., Ågerstrand, M., Rudén, C., 2009. WikiPharma – A freely available, easily accessible, interactive and comprehensive database for environmental effect data for pharmaceuticals. Regul. Toxicol. Pharm. 55, 367–371. 10.1016/j.yrtph.2009.08.009

Naeem, S., Wright, J.P., 2003. Disentangling biodiversity effects on ecosystem functioning: deriving solutions to a seemingly insurmountable problem. Ecol. Lett. 6, 567–579. 10.1046/j.1461-0248.2003.00471.x

Neale, P.A., Braun, G., Brack, W., Carmona, E., Gunold, R., König, M., Krauss, M., Liebmann, L., Liess, M., Link, M., Schäfer, R.B., Schlichting, R., Schreiner, V.C., Schulze, T., Vormeier, P., Weisner, O., Escher, B.I., 2020. Assessing the mixture effects in in vitro bioassays of chemicals occurring in small agricultural streams during rain events. Environ. Sci. Technol. 54, 8280–8290. 10.1021/acs.est.0c02235

Newman, M.C., 2009. Fundamentals of Ecotoxicology. CRC Press.

Nie, X., Gu, J., Lu, J., Pan, W., Yang, Y., 2009. Effects of norfloxacin and butylated hydroxyanisole on the freshwater microalga Scenedesmus obliquus. Ecotoxicology 18, 677–684. 10.1007/s10646-009-0334-1

Peters, K., Bundschuh, M., Schäfer, R.B., 2013. Review on the effects of toxicants on freshwater ecosystem functions. Environ. Pollut. 180, 324–329. 10.1016/j.envpol.2013.05.025

Poisot, T., 2011. The digitize package: extracting numerical data from scatterplots. The R Journal 3, 25–26.

R Core Team, 2024. R: a language and environment for statistical computing.

Ragnarsdottir, K.V., 2000. Environmental fate and toxicology of organophosphate pesticides. J. Geol. Soc. 157, 859–876. 10.1144/jgs.157.4.859

Reid, A.J., Carlson, A.K., Creed, I.F., Eliason, E.J., Gell, P.A., Johnson, P.T.J., Kidd, K.A., MacCormack, T.J., Olden, J.D., Ormerod, S.J., Smol, J.P., Taylor, W.W., Tockner, K., Vermaire, J.C., Dudgeon, D., Cooke, S.J., 2019. Emerging threats and persistent conservation challenges for freshwater biodiversity. Biol. Rev. 94, 849–873. 10.1111/brv.12480

Richmond, E.K., Rosi, E.J., Reisinger, A.J., Hanrahan, B.R., Thompson, R.M., Grace, M.R., 2019. Influences of the antidepressant fluoxetine on stream ecosystem function and aquatic insect emergence at environmentally realistic concentrations. J. Freshw. Ecol. 34, 513–531. 10.1080/02705060.2019.1629546

Robson, S.V., Rosi, E.J., Richmond, E.K., Grace, M.R., 2020. Environmental concentrations of pharmaceuticals alter metabolism, denitrification, and diatom assemblages in artificial streams. Freshw. Sci. 39, 256–267. 10.1086/708893

Rosi-Marshall, E.J., Royer, T.V., 2012. Pharmaceutical compounds and ecosystem function: an emerging research challenge for aquatic ecologists. Ecosystems 15, 867–880. 10.1007/s10021-012-9553-z

Rumschlag, S.L., Halstead, N.T., Hoverman, J.T., Raffel, T.R., Carrick, H.J., Hudson, P.J., Rohr, J.R., 2019. Effects of pesticides on exposure and susceptibility to parasites can be generalised to pesticide class and type in aquatic communities. Ecology Letters 22, 962–972. 10.1111/ele.13253

Rumschlag, S.L., Mahon, M.B., Hoverman, J.T., Raffel, T.R., Carrick, H.J., Hudson, P.J., Rohr, J.R., 2020. Consistent effects of pesticides on community structure and ecosystem function in freshwater systems. Nature Communications 11, 6333. 10.1038/s41467-020-20192-2

Schäfer, R.B., Bundschuh, M., Rouch, D.A., Szöcs, E., von der Ohe, P.C., Pettigrove, V., Schulz, R., Nugegoda, D., Kefford, B.J., 2012a. Effects of pesticide toxicity, salinity and other environmental variables on selected ecosystem functions in streams and the relevance for ecosystem services. Sci. Total Environ. 415, 69–78. 10.1016/j.scitotenv.2011.05.063

Schäfer, R.B., von der Ohe, P.C., Rasmussen, J., Kefford, B.J., Beketov, M.A., Schulz, R., Liess, M., 2012b. Thresholds for the effects of pesticides on invertebrate communities and leaf breakdown in stream ecosystems. Environ. Sci. Technol. 46, 5134–5142. 10.1021/es2039882

Schäfer, R.B., Gerner, N., Kefford, B.J., Rasmussen, J.J., Beketov, M.A., de Zwart, D., Liess, M., von der Ohe, P.C., 2013. How to Characterize Chemical Exposure to Predict Ecologic Effects on Aquatic Communities? Environ. Sci. Technol. 47, 7996–8004. 10.1021/es4014954

Schäfer, R.B., Baikova, D., Bayat, H., Beermann, A., Berger, S.A., Boenigk, J., Bruans, M., Burfeid-Castallanos, A., Cardinale, B., David, G., Feckler, A., Feld, C.K., Fink, P., Gessner, M.O., Hadziomerovic, U., Hering, D., le, T.T.Y., Macaulay, S., Madariaga, G.M., Mayombo, S., Pimentel, I.M., Orr, J., Oskapolor, S., Schlenker, A., Sures, B., Vermiert, A.-M., Weitere, M., Vos, M., Schürings, C., in preparation. Multiple stressors increase the effects of taxon loss on ecosystem functioning.

Scharmüller, A., Schreiner, V.C., Schäfer, R.B., 2020. Standartox: standardizing toxicity data. Data 5, 46. 10.3390/data5020046

Schulz, R., Bub, S., Petschick, L.L., Stehle, S., Wolfram, J., 2021. Applied pesticide toxicity shifts toward plants and invertebrates, even in GM crops. Science 372, 81–84. 10.1126/science.abe1148

Shiklomanov, I.A., 1993. World freshwater resource. Water in crisis: a guide to the world’s freshwater resources. Oxford University Press, New York.

Sinclair, T., Craig, P., Maltby, L.L., 2024. Climate warming shifts riverine macroinvertebrate communities to be more sensitive to chemical pollutants. Global Change Biology 30, e17254. 10.1111/gcb.17254

Soares, M.P., Jesus, F., Almeida, A.R., Domingues, I., Hayd, L., Soares, A.M.V.M., 2020. Effects of pH and nitrites on the toxicity of a cypermetrin-based pesticide to shrimps. Chemosphere 241, 125089. 10.1016/j.chemosphere.2019.125089

Stehle, S., Schulz, R., 2015. Agricultural insecticides threaten surface waters at the global scale. PNAS 112, 5750–5755. 10.1073/pnas.1500232112

Strayer, D.L., Dudgeon, D., 2010. Freshwater biodiversity conservation: recent progress and future challenges. J. N. Am. Benthol. Soc. 29, 344–358. 10.1899/08-171.1

Tilman, D., Isbell, F., Cowles, J.M., 2014. Biodiversity and ecosystem functioning. Annu. Rev. Ecol. Evol. Syst. 45, 471–493. 10.1146/annurev-ecolsys-120213-091917

Udy, J.W., Fellows, C.S., Bartkow, M.E., Bunn, S.E., Clapcott, J.E., Harch, B.D., 2006. Measures of nutrient processes as indicators of stream ecosystem health. Hydrobiologia 572, 89–102. 10.1007/s10750-005-9006-1

van Aert, R.C.M., Jackson, D., 2019. A new justification of the Hartung-Knapp method for random-effects meta-analysis based on weighted least squares regression. Res. Synth. Meth. 10, 515–527. 10.1002/jrsm.1356

Verheyen, J., Delnat, V., Theys, C., 2022. Daily temperature fluctuation can magnify the toxicity of pestides. Curr. Opin. Insect Sci. 51, 100919. 10.1016/j.cois.2022.100919

Viechtbauer, W., 2010. Conducting meta-analyses in R with the metafor package. J. Stat. Softw. 36, 1–48. 10.18637/jss.v036.i03

von Schiller, D., Acuña, V., Aristi, I., Arroita, M., Basaguren, A., Bellin, A., Boyero, L., Butturini, A., Ginebreda, A., Kalogianni, E., Larrañaga, A., Majone, B., Martínez, A., Monroy, S., Muñoz, I., Paunović, M., Pereda, O., Petrovic, M., Pozo, J., Rodríguez-Mozaz, S., Rivas, D., Sabater, S., Sabater, F., Skoulikidis, N., Solagaistua, L., Vardakas, L., Elosegi, A., 2017. River ecosystem processes: a synthesis of approaches, criteria of use and sensitivity to environmental stressors. Sci. Total Environ. 596–597, 465–480. 10.1016/j.scitotenv.2017.04.081

Wang, Z., Meador, J.P., Leung, K.M.Y., 2016. Metal toxicity to freshwater organisms as a function of pH: a meta-analysis. Chemosphere 144, 1544–1552. 10.1016/j.chemosphere.2015.10.032

Wang, Z., Walker, G.W., Muir, D.C.G., Nagatani-Yoshida, K., 2020. Toward a global understanding of chemical pollution: a first comprehensive analysis of national and regional chemical inventories. Environ. Sci. Technol. 54, 2575–2584. 10.1021/acs.est.9b06379

Warne, M.S.J., Hawker, D.W., 1995. The number of components in a mixture determines whether synergistic and antagonistic or additive toxicity predominate - the funnel hypothesis. Ecotoxicol. Environ. Safe. 31, 23–28. 10.1006/eesa.1995.1039

Wolfram, J., Stehle, S., Bub, S., Petschick, L.L., Schulz, R., 2021. Water quality and ecological risks in European surface waters – Monitoring improves while water quality decreases. Environ. Int. 152, 106479. 10.1016/j.envint.2021.106479

Young, R.G., Matthaei, C.D., Townsend, C.R., 2008. Organic matter breakdown and ecosystem metabolism: functional indicators for assessing river ecosystem health. J. N. Am. Benthol. Soc. 27, 605–625. 10.1899/07-121.1

Zubrod, J.P., Bundschuh, M., Arts, G., Brühl, C.A., Imfeld, G., Knäbel, A., Payraudeau, S., Rasmussen, J.J., Rohr, J., Scharmüller, A., Smalling, K., Stehle, S., Schulz, R., Schäfer, R.B., 2019. Fungicides: an overlooked pesticide class? Environ. Sci. Technol. 53, 3347–3365. 10.1021/acs.est.8b04392

