## Supplementary material for "Meta-analysis on the effects of chemical stressors on freshwater ecosystem functions": Fig. A1

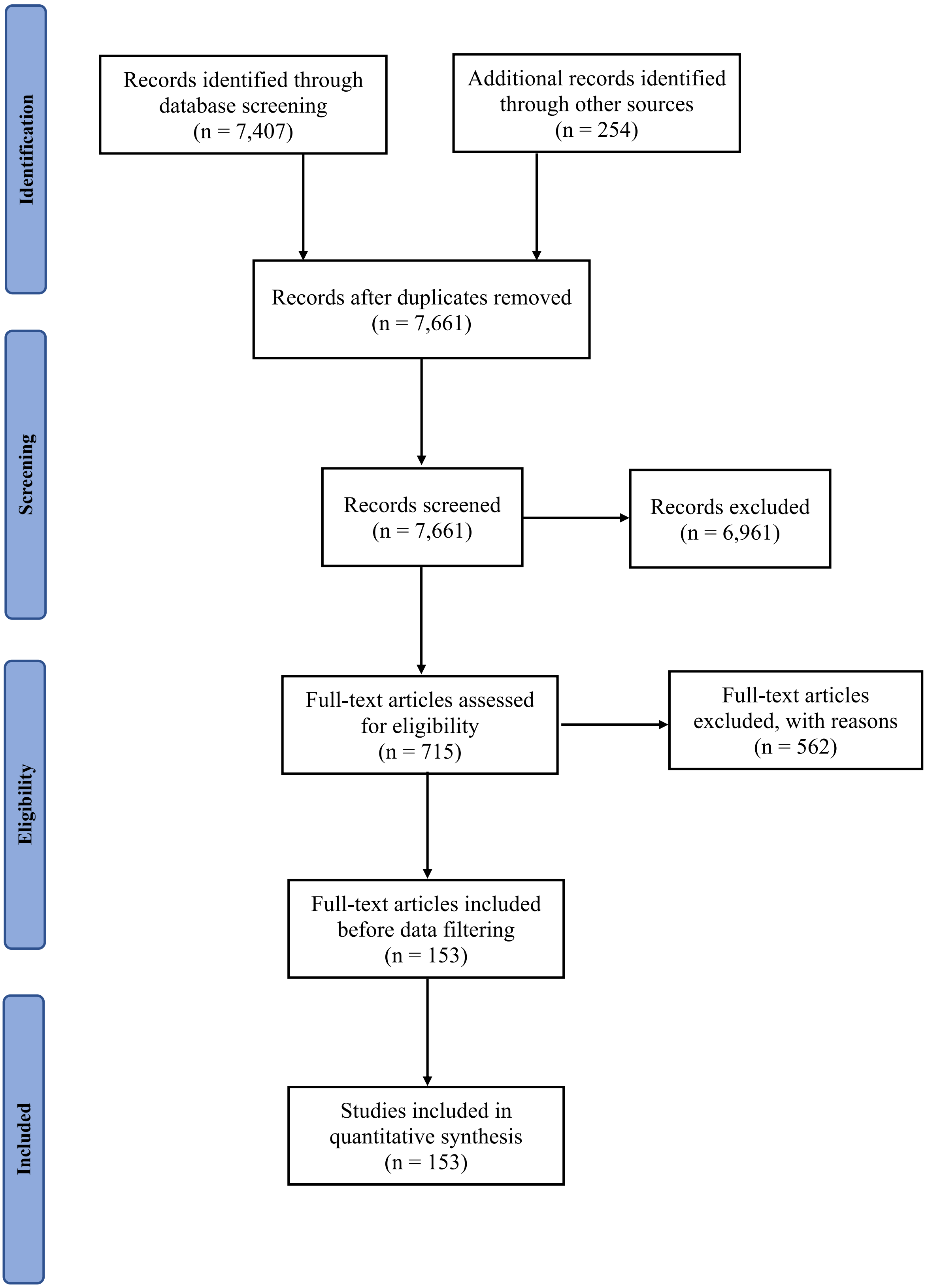


Figure A1: PRISMA (Preferred Reporting Items for Systematic Reviews and Meta-Analyses; Page et al. (2021)) flow chart illustrating how 7,661 initial records were narrowed down to 153 studies from which 350 effect sizes were extracted for final analyses.

Table A1: Overview of chemical stressors and respective concentrations required to obtain an effect in 50% of the exposed most sensitive species (EC50; mg/L; sorted by organism group) and sources of the data. If not further specified, the data were extracted from Standartox (Scharmüller et al. 2020), the Pesticide Property Database (Lewis et al. 2016), the Veterinary Substance Database (Lewis et al. 2011), the Norman Database (https://www.norman-network.com/nds/), and WikiPharma (Molander et al. 2009).

| **Chemical stressor** | **$EC_{50}$** | **Organism group** | **Species** | **Source** | **Note** |
|---|---|---|---|---|---|
| 2,4-D | 55 | Algae | *Gomphonema sp.* | Standartox | |
| Abacavir | 55.75 | Algae | *Selenastrum capricornutum* | Norman Database | Derived from PNEC & AF of 1000 |
| Aluminum | 1000000 | Algae | *Raphidocelis subcapitata* | Van Hoecke et al. (2011) | |
| Amoxicillin | 37 | Algae | *Microcystis aeruginosa* | VSDB | |
| Arsenic | 34700 | Algae | *R. subcapitata* | Tišler and Zagorc-Končan (2002) | |
| Atrazine | 12.9614814 | Algae | *Pseudoanabaena galeata* | Standartox | |
| Atrazine | 0.029 | Aquatic plants | *Stuckenia pectinata* | Standartox | |
| Azoxystrobin | 106 | Algae | *R. subcapitata* | Standartox | |
| Azoxystrobin | 56 | Aquatic invertebrates | *Americamysis.bahia* | Standartox | |
| Bensulfuron-methyl | 20 | Algae | *R. subcapitata* | PPDB | |
| Boscalid | 1340 | Algae | *R. subcapitata* | Standartox | |
| Butachlor | 9.64443663 | Algae | *R. subcapitata* | Standartox | |
| Cadmium | 109 | Algae | *S. capricornutum* | van der Heever and Grobbelaar (1995) | |
| Cadmium | 101 | Aquatic invertebrates | *Daphnia magna* | Shaw et al. (2006) | |
| Cadmium | 323 | Aquatic plants | *Lemna minor* | Naumann et al. (2007) | |
| Carbamazepine | 2275.78 | Algae | *S. capricornutum* | Norman Database | Derived from PNEC & AF of 1000 |
| Carbendazim | 340 | Algae | *Chlorella pyrenoidosa* | Standartox | |
| Carbendazim | 43.44331 | Aquatic invertebrates | *Gammarus pulex* | Standartox | |
| Chlorantraniliprole | 4 | Aquatic invertebrates | *Chironomus dilutus* | Standartox | |

| | | | | | |
|---|---|---|---|---|---|
| Chlorothalonil | 2 | Algae | *C. pyrenoidosa* | Standartox | |
| Chlorothalonil | 0.97 | Aquatic invertebrates | *Dreissena polymorpha* | Standartox | |
| Chlorpyrifos | 38 | Algae | *Nitzschia closterium* | Standartox | |
| Chromium | 970 | Algae | *R. subcapitata* | Paixão et al. (2008) | |
| Ciprofloxacin | 4830 | Algae | *R. subcapitata* | WikiPharma | |
| Clotrimazole | 29.98 | Algae | *S. capricornutum* | Norman Database | Derived from PNEC & AF of 1000 |
| Copper | 552 | Algae | *S. capricornutum* | van der Heever and Grobbelaar (1995) | |
| Copper | 950.5 | Aquatic invertebrates | *D. magna* | Malaj et al. (2012) | |
| Copper | 330 | Aquatic plants | *L. minor* | Naumann et al. (2007) | |
| Copper sulfate | 2300 | Aquatic invertebrates | *D. magna* | PPDB | |
| Copper-hydroxide | 38 | Aquatic invertebrates | *D. magna* | PPDB | |
| Cu-hydroxide | 9 | Algae | *R. subcapitata* | PPDB | |
| Cu-sulphate | 12300 | Algae | *R. subcapitata* | PPDB | |
| Cypermethrin | 62.6921454 | Algae | *Skeletonema costatum* | Standartox | |
| Cyprodinil | 1970 | Algae | *S. costatum* | Standartox | |
| Cyprodinil | 32 | Aquatic invertebrates | *D. magna* | Standartox | |
| Diazinone | 0.03326532 | Aquatic invertebrates | *D. magna* | Standartox | |
| Diclofenac | 21300 | Algae | *R. subcapitata* | WikiPharma | |
| Dimethoate | 5500 | Algae | *Chlamydomonas noctigama* | Standartox | |
| Diuron | 1.1880867 | Algae | *Chara vulgaris* | Standartox | |
| Diuron | 5 | Aquatic plants | *Myriophyllum spicatum* | Standartox | |
| Epoxiconazole | 1.32E+03 | Aquatic invertebrates | *C. riparius* | Standartox | |
| Erythromycin | 20 | Algae | *R. subcapitata* | VSDB | |
| Etofenprox | 0.0188 | Aquatic invertebrates | *A. bahia* | Standartox | |

| | | | | | |
|---|---|---|---|---|---|
| Fexofenadine | 53.44 | Algae | *S. capricornutum* | Norman Database | Derived from PNEC & AF of 1000 |
| Fludioxonil | 24 | Algae | *Scendesmus subspicatus* | PPDB | |
| Fludioxonil | 920 | Aquatic plants | *L. gibba* | PPDB | |
| Flumioxazin | 0.35 | Aquatic plants | *L. gibba* | PPDB | |
| Fluoxetine | 0.000052 | Algae | *Skeletonema costatum* | WikiPharma | |
| Fluridone | 17 | Algae | *Thalassiosira weissflogii* | Standartox | |
| Glyphosate | 5.3 | Algae | *Gomphonema* | Standartox | |
| Glyphosate | 12000 | Aquatic plants | *L. gibba* | PPDB | |
| Ibuprofen | 315000 | Algae | *Desmodesmus subspicatus* | WikiPharma | |
| Imazalil | 3500 | Aquatic invertebrates | *D. magna* | PPDB | |
| Imidacloprid | 212424.1 | Algae | *D. subspicatus* | Standartox | |
| Imidacloprid | 85000 | Aquatic invertebrates | *D. magna* | PPDB | |
| Isoproturon | 5 | Algae | *C. pyrenoidosa* | Standartox | |
| lambda-cyhalothrin | 0.00382921 | Aquatic invertebrates | *Hyalella azteca* | Standartox | |
| Lead | 10.35 | Algae | *R. subcapitata* | Chen and Lin (1997) | |
| Levofloxacin | 1200 | Algae | *R. subcapitata* | VSDB | |
| Lindane | 2.771281 | Aquatic invertebrates | *Limnephilus lunatus* | Standartox | |
| Linuron | 16 | Algae | *R. subcapitata* | PPDB | |
| Linuron | 0.07 | Aquatic plants | *S. pectinata* | Standartox | |
| Manganese | 5200 | Algae | *S. quadricauda* | Fargašová et al. (1997) | |
| Mercury | 78.5 | Algae | *S. capricornutum* | van der Heever and Grobbelaar (1995) | |
| Nickel | 143 | Algae | *R. subcapitata* | Chen and Lin (1997) | |
| Norfloxacin | 16600 | Algae | *S. capricornutum* | VSDB | |
| Norflurazon | 9.7 | Algae | *R. subcapitata* | Standartox | |
| Paraquat | 0.23 | Algae | *R. subcapitata* | PPDB | |

| | | | | | |
|---|---|---|---|---|---|
| Procymidone | 2600 | Algae | *Scenedesmus acutus* | PPDB | |
| Propriconazole | 510 | Aquatic invertebrates | *A. bahia* | Standartox | |
| Pyraclostrobin | 4.16 | Aquatic invertebrates | *A. bahia* | Standartox | |
| Pyrimethanil | 1200 | Algae | *R. subcapitata* | PPDB | |
| Pyrimethanil | 7800 | Aquatic plants | *L. gibba* | PPDB | |
| Quinoxyfen | 27 | Algae | *R. subcapitata* | Standartox | |
| Quinoxyfen | 72 | Aquatic invertebrates | *Crassostrea virginica* | Standartox | |
| s-Metalaxyl | 420 | Algae | *S. subcapitatus* | PPDB | |
| S-metolachlor | 17 | Algae | *R. subcapitata* | PPDB | |
| Silver | 0.29 | Algae | *R. subcapitata* | Mertens et al. (2019) | |
| Simazine | 24.4 | Algae | *Nephroselmis pyriformis* | Standartox | |
| Simazine | 104.73956 | Aquatic plants | *Vallisneria americana* | Standartox | |
| Sulfamethoxazole | 2743.92 | Algae | *S. capricornutum* | Norman Database | Derived from PNEC & AF of 1000 |
| Tebuconazole | 1450 | Algae | *D. subspicatus* | Standartox | |
| Tebuconazole | 4.863127 | Aquatic invertebrates | *D. galeata* | Standartox | |
| Thiacloprid | 1.06 | Aquatic invertebrates | *C. tepperi* | Standartox | |
| Thiobencarb | 17 | Algae | *R. subcapitata* | PPDB | |
| Trichlorofon | 10000 | Algae | *S. subcapitatus* | VSDB | |
| Zinc | 138 | Algae | *R. subcapitata* | Muyssen and Janssen (2001) | |
| Zinc | 909 | Aquatic plants | *L. minor* | Naumann et al. (2007) | |
| Zineb | 232.249 | Algae | *N. pungens* | Standartox | |

### References

Chen C-Y, Lin K-C. 1997. Optimization and performance evaluation of the continuous algal toxicity test. Environ. Toxicol. Chem. 16, 1337–1344. doi:10.1002/etc.5620160701

Fargašová A, Bumbálová A, Havránek E. 1997. Metal bioaccumulation by the freshwater alga *Scenedesmus quadricauda*. J. Radioanal. Nucl. Chem. 218, 107–110. doi:10.1007/BF02033984

Lewis K, Tzilivakis J, Green A, Warner DJ. 2011. Veterinary Substances Database (VSDB).

Lewis KA, Tzilivakis J, Warner DJ, Green A. 2016. An international database for pesticide risk assessments and management. Hum. Ecol. Risk Assess. 22, 1050–1064. doi:10.1080/10807039.2015.1133242

Malaj E, Ohe PC von der, Grote M, Kühne R, Mondy CP, Usseglio-Polatera P, Brack W, Schäfer RB. 2014. Organic chemicals jeopardize the health of freshwater ecosystems on the continental scale. PNAS 111:9549–9554. doi:10.1073/pnas.1321082111

Mertens J, Oorts K, Leverett D, Arijs K. 2019. Effects of silver nitrate are a conservative estimate for the effects of silver nanoparticles on algae growth and *Daphnia magna* reproduction. Environ. Toxicol. Chem. 38, 1701–1713. doi:10.1002/etc.4463

Molander L, Ågerstrand M, Rudén C. 2009. WikiPharma – A freely available, easily accessible, interactive and comprehensive database for environmental effect data for pharmaceuticals. Regul. Toxicol. Pharm. 55, 367–371. doi:10.1016/j.yrtph.2009.08.009

Muyssen BTA, Janssen CR. 2001. Zinc acclimation and its effect on the zinc tolerance of *Raphidocelis subcapitata* and *Chlorella vulgaris* in laboratory experiments. Chemosphere 45, 507–514. doi:10.1016/S0045-6535(01)00047-9

Naumann B, Eberius M, Appenroth K-J. 2007. Growth rate based dose–response relationships and EC-values of ten heavy metals using the duckweed growth inhibition test (ISO 20079) with *Lemna minor* L. clone St. J. Plant Physiol. 164, 1656–1664. doi:10.1016/j.jplph.2006.10.011

Page MJ, McKenzie JE, Bossuyt PM, Boutron I, Hoffmann TC, Mulrow CD, Shamseer L, Tetzlaff JM, Akl EA, Brennan SE, Chou R, Glanville J, Grimshaw JM, Hróbjartsson A, Lalu MM, Li T, Loder EW, Mayo-Wilson E, McDonald S, McGuinness LA, Stewart LA, Thomas J, Tricco AC, Welch VA, Whiting P, Moher D. 2021. The PRISMA 2020 statement: an updated guideline for reporting systematic reviews. Int. J. Surg. 88, 105906. doi:10.1016/j.ijsu.2021.105906

Paixão SM, Silva L, Fernandes A, O'Rourke K, Mendonça E, Picado A. 2008. Performance of a miniaturized algal bioassay in phytotoxicity screening. Ecotoxicology 17, 165–171. doi:10.1007/s10646-007-0179-4

Scharmüller A, Schreiner VC, Schäfer RB. 2020. Standartox: standardizing toxicity data. Data 5, 46. doi:10.3390/data5020046

Shaw JR, Dempsey TD, Chen CY, Hamilton JW, Folt CL. 2006. Comparative toxicity of cadmium, zinc, and mixtures of cadmium and zinc to daphnids. Environ. Toxicol. Chem. 25, 182–189. doi:10.1897/05-243R.1

Tišler T, Zagorc-Končan J. 2002. Acute and chronic toxicity of arsenic to some aquatic organisms. Bull. Environ. Contam. Toxicol. 69, 421–429. doi:10.1007/s00128-002-0079-5

van der Heever JU, Grobbelaar JA. 1995. The use of *Selenastrum capricornutum* growth potential as a measure of toxicity of a few selected compounds. Water SA 22, 183–191.

Van Hoecke K, De Schamphelaere KAC, Ramirez–Garcia S, Van der Meeren P, Smagghe G, Janssen CR. 2011. Influence of alumina coating on characteristics and effects of SiO2 nanoparticles in algal growth inhibition assays at various pH and organic matter contents. Environ. Int. 37, 1118–1125. doi:10.1016/j.envint.2011.02.009
